# National-scale biogeography and function of river and stream bacterial biofilm communities

**DOI:** 10.1101/2025.03.05.641783

**Authors:** Amy C. Thorpe, Susheel Bhanu Busi, Jonathan Warren, Laura H. Hunt, Kerry Walsh, Daniel S. Read

## Abstract

Biofilm-dwelling microorganisms coat the surfaces of stones in river and stream ecosystems, forming diverse communities that are fundamental to biogeochemical processes and ecosystem functioning^1,2^. Flowing water (lotic) ecosystems are under pressure from a wide range of interacting stressors including changes in land use, chemical pollution, and climate^3^. Despite their ecological importance, the taxonomic and functional diversity of river biofilms and their responses to environmental change are limited by a lack of understanding of their taxonomic composition and physicochemical drivers across large spatial scales. We conducted a national-scale assessment of bacterial diversity and function using metagenomic sequencing from rivers and streams across England, analogous to other large-scale efforts to understand microbial biogeography across diverse environments^4,5,6,7^. We recovered 1,014 metagenome-assembled genomes (MAGs) from 450 biofilms collected across England’s extensive river network, revealing substantial taxonomic novelty, with ∼20% of the MAGs representing novel genera. We demonstrated that biofilm communities, dominated by generalist bacteria, exhibit remarkable functional diversity and metabolic versatility, and play a significant role in nutrient cycling with the potential for contaminant transformation. Environmental drivers, most notably geology, land cover, and nutrients, explained up to 90% of the variation in community composition. These findings highlight the importance of river biofilms and establish a foundation for future research on the roles of biofilms in ecosystem health and resilience to environmental change.

## Introduction

It is estimated that up to 80% of all bacterial and archaeal cells on Earth exist within biofilms^8^. These biofilms dominate microbial life in freshwater streams and rivers^2^, hosting complex, interacting assemblages of taxonomically and functionally diverse heterotrophic and phototrophic microbes embedded within an extracellular polymeric matrix that adheres to the surfaces of stones, plants, and sediments^1^. Biofilms are hotspots of metabolic activity, driving essential biogeochemical processes such as nutrient cycling, primary production, and respiration^9^, degrading pollutants, regulating water quality^10^, and supporting energy flow to higher trophic levels^11^. Microbes within biofilms are critical to the functioning of river ecosystems^1^, which are under increasing pressure from pollution and climate change^3^. Understanding the diversity and function of biofilm microbial communities in rivers is, therefore, essential for managing and conserving these vital ecosystems^10^.

Diverse microbial communities have been detected in river biofilms^1,2^. The composition and functionality of these communities are likely influenced by a complex interplay of factors, including physiochemical parameters such as temperature, pH, and dissolved oxygen^12,13,14^; the concentrations of nutrients, organic matter, and pollutants^15,16,17^; flow dynamics^18^; land use^19,20^; and processes such as selection and dispersal^21^. By shaping microbial community composition and function, environmental changes can significantly affect biogeochemical fluxes and overall ecosystem functions^1^. However, our understanding of the diversity of microbial communities and their functional roles in river biofilms is limited. Furthermore, identifying the environmental drivers influencing biofilm community dynamics and assessing the resilience of these communities to pressures, such as nutrient pollution, presents an ongoing challenge.

Many studies of river biofilm communities have focused on individual rivers or catchments, providing valuable insights into local environmental drivers of microbial diversity and community dynamics^22,23,24^. However, these studies typically comprise a relatively small number of samples and cover a limited geographic area. In recent years, national-scale studies have been undertaken to capture the wider diversity of freshwater microbes and explore their biogeographic patterns across a broad range of environments^4,5,6,25^. These national-scale datasets demonstrate the value of molecular approaches, including metagenomics, for uncovering both the taxonomic and functional diversity of microbial communities. Moreover, they highlight community dynamics and adaptations in response to environmental drivers across large spatial scales. Collectively, national-scale datasets can provide a more comprehensive understanding of the ecological dynamics of freshwater microbes and their drivers at a global scale. However, there are few examples of such national-scale studies of river biofilm communities, and, to our knowledge, no national-scale metagenomic assessment of river microbes currently exists in England.

### National River Surveillance Network (RSN)

The Environment Agency (EA) established the River Surveillance Network (RSN) as a national initiative to monitor and assess long-term changes in the health of English rivers. The RSN comprises over 2,600 sites, representing the diversity of England’s extensive river network. As part of this initiative, 450 river biofilms were systematically collected over a three-year period from 146 RSN sites (Fig. 1A). This comprehensive dataset spanned a latitudinal gradient of 645 km. It encompassed all major land cover types present in the country, ranging from woodlands and grasslands to arable land and urban areas, and represented a variety of catchment geologies (Extended Data Table 1, Supplementary Data 1). In addition to the diverse catchments, the RSN sites spanned considerable variability in their physicochemical conditions, including water temperature, alkalinity, pH, dissolved oxygen, dissolved organic carbon, orthophosphate, nitrate-N, and nitrite-N (Extended Data Table 2, Supplementary Data 1).

**Fig. 1.**
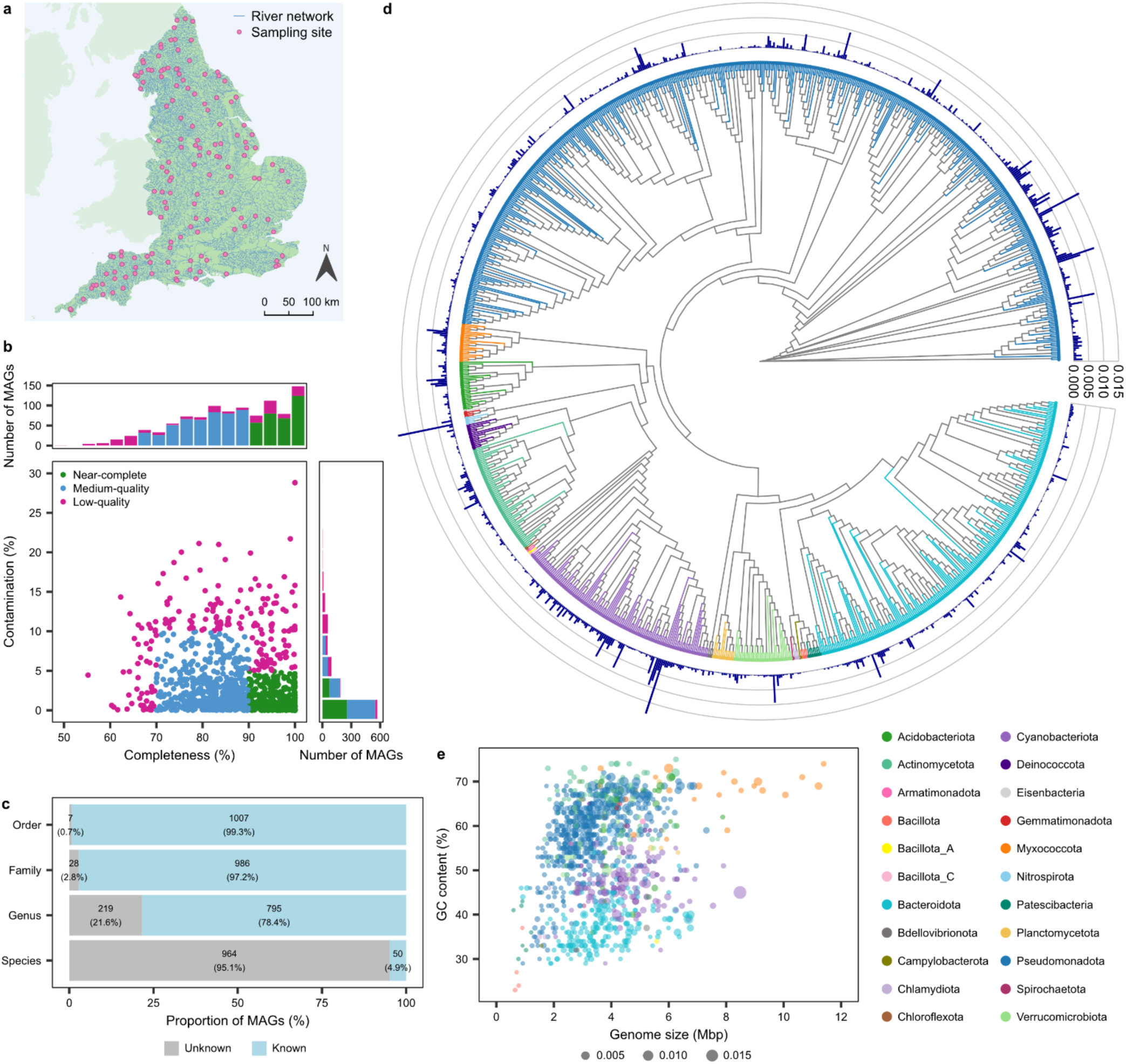
Overview of the diversity, novelty, and genomic characteristics of river biofilm bacterial MAGs. (**a)** Biofilm sampling sites in rivers across England (contains OS data © Crown copyright and database rights 2024). **(b)** Distribution of MAG completeness and contamination. **(c)** Taxonomic novelty of MAGs at the order, family, genus, and species level. **(d)** Phylogenetic tree of MAGs where colours represent different bacterial phyla, and outer bars represent the mean relative abundance of MAGs. **(e)** Estimated GC content and genome size of MAGs where point size is scaled to mean relative abundance.

This systematic investigation of the river network, coupled with detailed environmental data, enables a robust assessment of microbial biofilms, comparable to existing large-scale efforts^4,5,6,7^, capturing their biogeographic distribution, taxonomic diversity, functional capabilities, and environmental drivers.

### Diversity and composition of river biofilm microbial communities

River biofilms are cosmopolitan in their distribution and known to host high microbial diversity, supported by continuous nutrient and community inputs from upstream^1^. However, our understanding of freshwater biofilm microbial communities is significantly less advanced than that of microbial communities in the water column or sediments^12^. Although several studies have explored the taxonomic composition of river biofilms at the local scale^13,14,23^, the overall abundance, composition, and genomic traits of these communities across large spatial scales remain poorly understood.

Using metagenomic data averaging 66.80 million reads per sample (± 28.5 million reads SD), we found that bacterial sequences comprised the majority (85.17%) of all metagenomic reads in the river biofilms. Eukaryotes and Archaea represented 11.56% and 2.64% of the reads, respectively (Supplementary Data 2). Because of their dominance in biofilms, we focused our analysis on bacterial communities. The assembled reads were used to reconstruct a total of 4,027 bacterial bins (archaeal bins were not recovered), which were dereplicated into 1,014 metagenome-assembled genomes (MAGs) (also known as species-level genome bins). Of these, 329 MAGs (32%) were identified as near-complete, 491 (48%) as medium-quality, and 194 (19%) as low-quality (Fig. 1B) with a mean completeness of 86.70% (± 9.3% SD) and a mean contamination of 3.57% (± 4.1% SD). Mean genome size was estimated to be 3.66 Mbp (± 1.4 Mbp SD) and mean GC content was 52.66% (± 11.7% SD) (Extended Data Fig. 1, Supplementary Data 3).

The 1,014 bacterial MAGs encompassed a diverse range of taxa, including 22 known phyla, 38 classes, 96 orders, 173 families, 339 genera and 51 species, and represented a substantial taxonomic novelty, with 219 (21.6%) of the recovered MAGs representing previously uncharacterised genera, and 964 (95.1%) representing previously uncharacterised species (Fig. 1C). Stringent assessment using only near-complete MAGs revealed that 306 (93%) were novel species-level genome bins, with no previously identified representatives. Across all river biofilms, Pseudomonadota was the most dominant phylum and comprised almost half of the total community, with a mean relative abundance of 48.08% (± 14.8% SD) (Fig. 1D). Other abundant phyla included Cyanobacteriota with a mean relative abundance of 18.12% (± 17.9%), Bacteroidota with a mean relative abundance of 13.72% (± 9.1%), and Actinomycetota with a mean relative abundance of 5.61% (± 4.6%). Other less abundant phyla each comprised less than 5% of the total community on average. This community composition aligns with previous studies on biofilms collected from rivers^26,27^ and glacier-fed streams^28^, which also reported the dominance of Pseudomonadota and Bacteroidota, which have been reported to play important roles in the degradation of organic matter, and Cyanobacteriota, which contributes to primary productivity^29^.

We found that genome size and GC content were correlated with phylogenetic background, delineating these traits across the bacterial tree of life (Fig. 1E). It was observed that Myxococcota have large genomes with a high GC content, Pseudomonadota have small to medium genomes with a high GC content, and Cyanobacteriota and Bacteroidota have small to medium genomes with a lower GC content. A similar phylogenetic relationship between GC content and genome size has been observed in lake bacteria^30,31^. These traits may reflect ecological niche adaptation. For example, bacteria with larger genomes are typically copiotrophs, with numerous genes associated with complex metabolic processes, allowing them to thrive in nutrient-rich environments. In contrast, those with smaller genomes often have specific nutrient requirements and occupy more stable, oligotrophic environments^32^. Furthermore, lower GC content reduces nitrogen requirements, which may be advantageous for bacteria in nitrogen-limited environments^31^. To test this hypothesis, we assessed the relationship between measured nutrient concentrations (nitrate-N, nitrite-N, and orthophosphate) and genome size, which did not reveal any strong association (R <0.002, data not shown). Genome size distribution is likely affected by a complex interplay of environmental factors, in addition to trophic strategies^33^.

### Biogeography of river biofilm bacteria

River biofilms host complex communities composed of both generalist and specialist taxa, each fulfilling distinct ecological roles^2^. Generalist taxa are highly adaptable, thriving across a diversity of habitats, and tolerating a broad range of environmental conditions, whereas specialist taxa are adapted to specific habitats, thriving only under specific environmental conditions^34^. Despite the ecological significance of these communities, the biogeographic patterns of river biofilm bacteria are poorly understood and have not yet been addressed in the context of ecological niche space. Using biogeographic mapping and niche breadth analysis coupled with measures of abundance and occupancy, we investigated the spatial distribution and ecological niches of river biofilm bacteria.

The distribution of MAGs across the river network revealed national-scale biogeographic patterns at the phylum level (Fig. 2). Pseudomonadota and Bacteroidota exhibited high relative abundances throughout the country (>5% in more than 90% of sites), dominating biofilm communities. In contrast, Cyanobacteriota displayed regional preferences, with higher relative abundance in the north and southwest of England, and a lower occupancy overall (Fig. 3A), indicating that they occasionally dominate biofilm communities. Other taxa, such as Acidobacteriota, Myxococcota, and Nitrospirota, had a widespread presence but exhibited localised hotspots where their relative abundance was notably higher. Campylobacter and Chloroflexota had more restricted biogeographic distributions and were present at low relative abundances in only 10 and 79 samples, respectively. However, most MAGs are widely distributed rather than limited to specific locations. Each MAG occupied an average of 417 samples (93%) spatially distributed across the river network, with 227 MAGs (22%) present in all samples. Many of these 227 high-occupancy MAGs were the most abundant members of the community, with a mean relative abundance of up to 1.90% (Fig. 3A). A similar relationship between occupancy and mean relative abundance has been observed in surface water bacterial communities in rivers across North America^4^. However, while river networks comprise mostly high-occupancy taxa, studies of bacterial communities in spatially distributed lakes have found a greater proportion of low-occupancy taxa^5,35^. This difference likely reflects the underlying ecological factors that differ between rivers and lakes, such as habitat connectivity, dispersal dynamics, and environmental stability^36^.

**Fig. 2.**
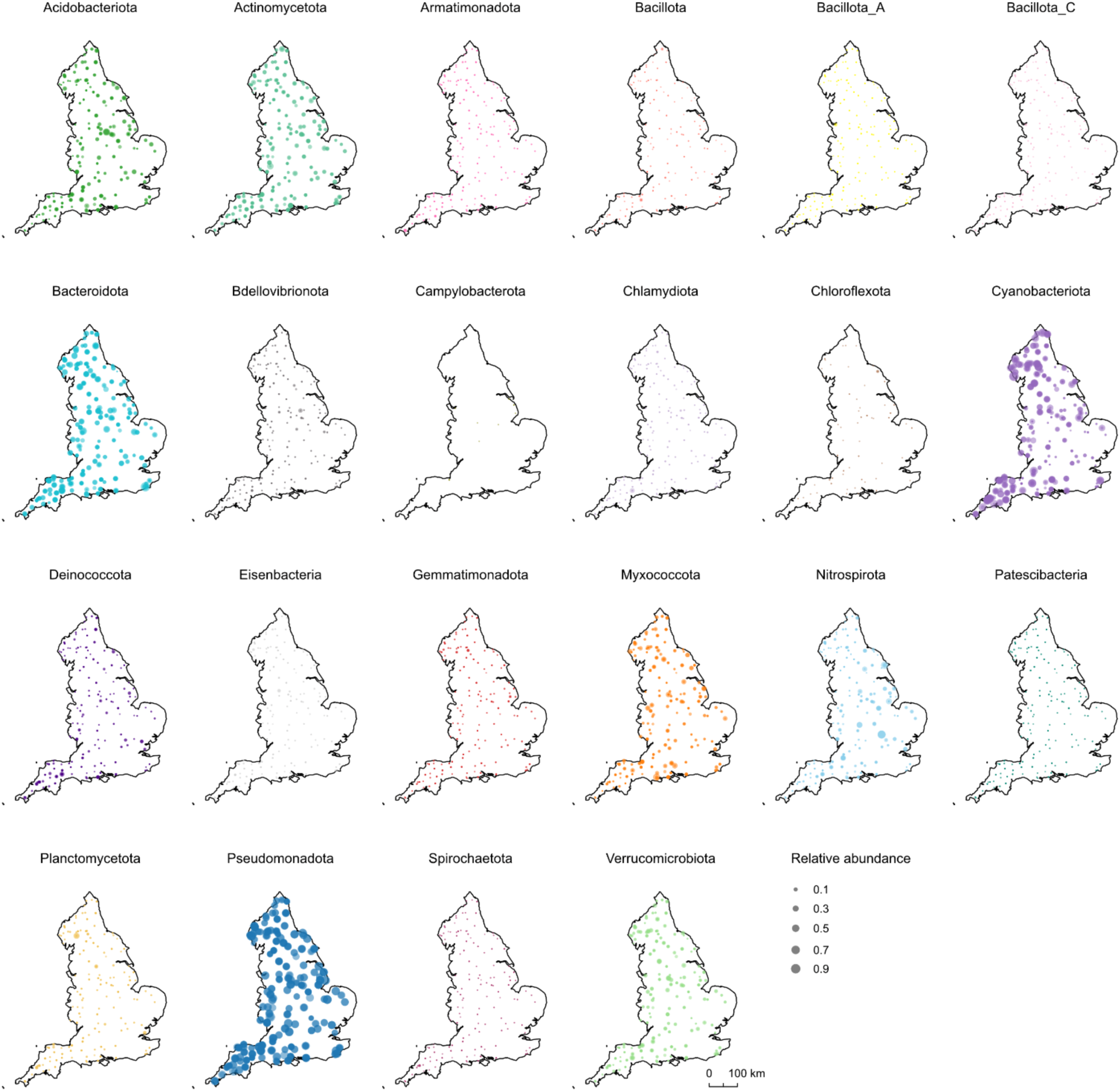
Biogeographic distribution of bacterial MAGs in rivers across England. Colours represent different bacterial phyla and points are scaled to relative abundance.

**Fig. 3.**
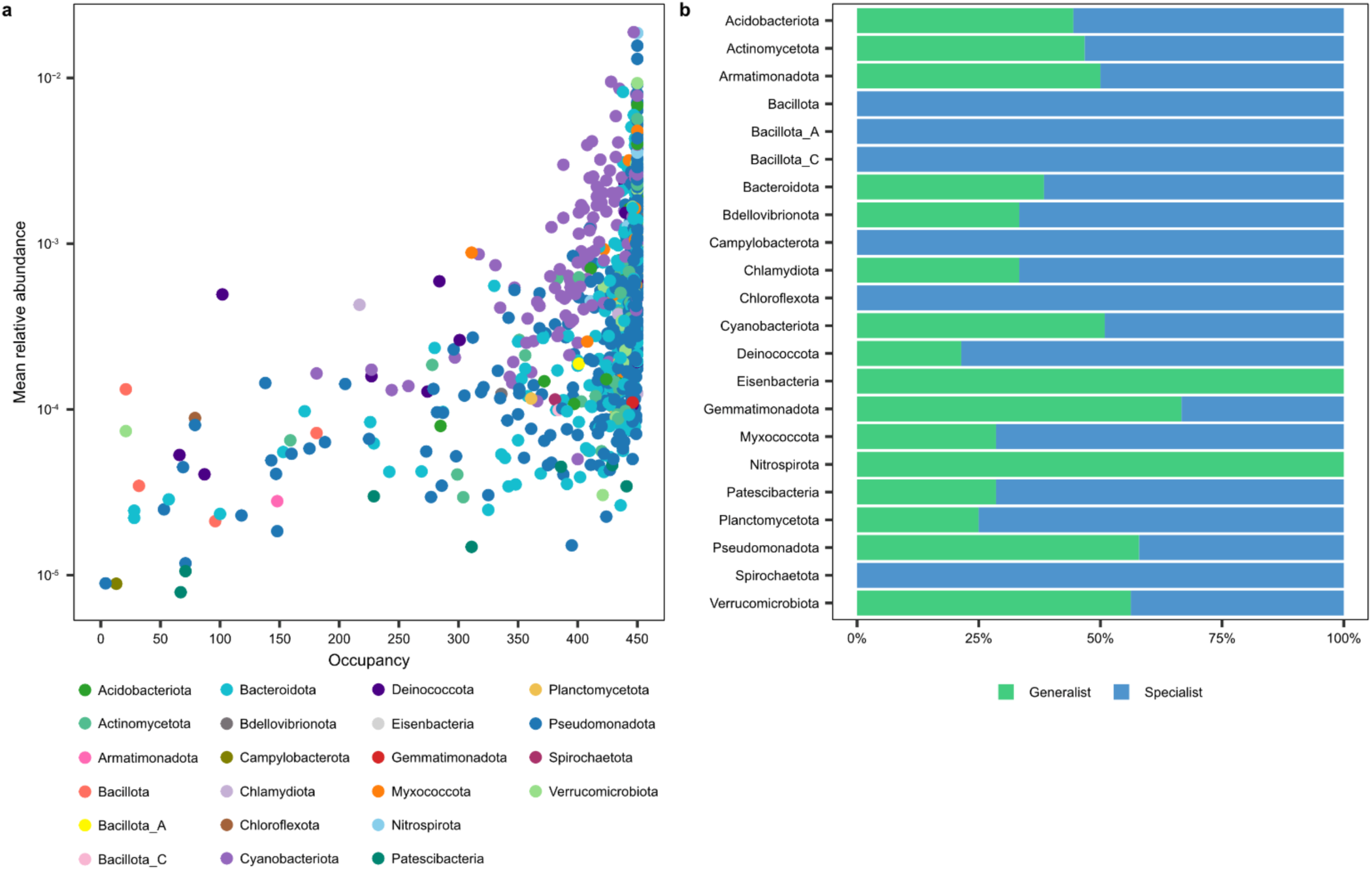
Ecological niche breadth of bacterial MAGs. **(a)** Occupancy and mean relative abundance (log scale) of MAGs where point colour represents bacterial phylum. **(b)** Proportion of MAGs of each bacterial phylum classified as generalists and specialists.

Using niche breadth analysis, we further categorised the MAGs as generalists (niche breadth, Bn > median Bn) or specialists (Bn < median Bn). Among the 227 high-occupancy MAGs, 71% were identified as generalists, and among the most abundant phyla, 57.95% of Pseudomonadota MAGs and 50.88% of Cyanobacteriota MAGs were categorised as generalists (Fig. 3B). This finding is supported by previous studies which have also detected a high abundance of generalist bacteria in river and stream biofilms^2,37,38^. Furthermore, the distance-decay relationship of bacterial community dissimilarity and geographic distance was positive, but weak (R^2^ = 0.04, p < 0.001) (Extended Data Fig. 2A and B), suggesting dispersal rates in bacterial biofilms is high, although some species sorting due to local environmental conditions likely occurs. The dominance of high-occupancy, generalist MAGs across the river network emphasises the role of rivers as highly interconnected ecosystems with potential for widespread microbial dispersal. The ability of generalist bacteria to tolerate a diverse range of environmental conditions is likely an advantage in lotic ecosystems, where biofilm communities may become dislodged and recolonise further downstream^37^.

### Metabolic and functional potential of river biofilm communities

Biofilm communities play critical roles in ecological processes such as primary production, nutrient cycling, and the transformation and breakdown of organic material and pollutants^9,39^. While their biogeochemical cycling contributions can strongly influence the health and productivity of the wider river ecosystem by mediating the availability of nutrients and other compounds to higher trophic levels in the food web^11^, our understanding of the functional diversity of river biofilms is limited. Studies have emphasised the importance of assessing the metabolic capabilities of biofilms to better understand their functional roles in river ecosystems^1^.

Several bacterial phyla contributed to carbon, nitrogen, and sulfur cycling in the river biofilms, and among these, Pseudomonadota, Cyanobacteriota, and Bacteroidota were the primary contributors, contributing more than 46%, 13% and 7% of genes involved in these cycles (Extended Data Fig. 3). Across all 1,014 MAGs, the majority of steps in carbon, nitrogen, and sulfur cycling pathways were represented (Fig. 4A-C). Nearly all MAGs encoded for genes involved in organic carbon oxidation (99.05%), and a substantial proportion of the MAGs included genes involved in fermentation (86.19%), acetate oxidation (72.78%), sulfur oxidation (47.44%), and nitrite ammonification (43.69%). Many of these pathways represented by the river biofilms, including organic carbon oxidation and sulfur oxidation, were also prevalent in mountain stream biofilms^40^. Carbon cycling genes, such as *sgdh* (pentose phosphate pathway) and *RuBisCO IV* (carbon fixation), were abundant in Pseudomonadota and Cyanobacteriota (Fig. 4D). Nitrogen cycling genes, including *nirD* and *nirB* (nitrite reductases) (Fig. 4E), and sulfur cycling genes, including *sdo* (sulfur dioxygenase), *fccB* (sulfur dehydrogenase) and *sat* (sulfate reduction) (Fig. 4F), were particularly abundant in Pseudomonadota, Cyanobacteriota, and Bacteroidota. Additionally, Pseudomonadota and Acidobacteriota encoded the most diverse range of genes involved in these pathways, with Pseudomonadota encoding 17, 16, and 10 genes and Acidobacteriota encoding 13, 15, and 9 genes involved in the carbon, nitrogen, and sulfur cycles, respectively. The biofilm MAGs also encoded a range of genes involved in oxygen and hydrogen cycling (Extended Data Fig. 4, Supplementary Data 4). These results demonstrate that biofilm communities have significant metabolic potential and capacity to perform diverse biochemical roles in river ecosystems.

**Fig. 4.**
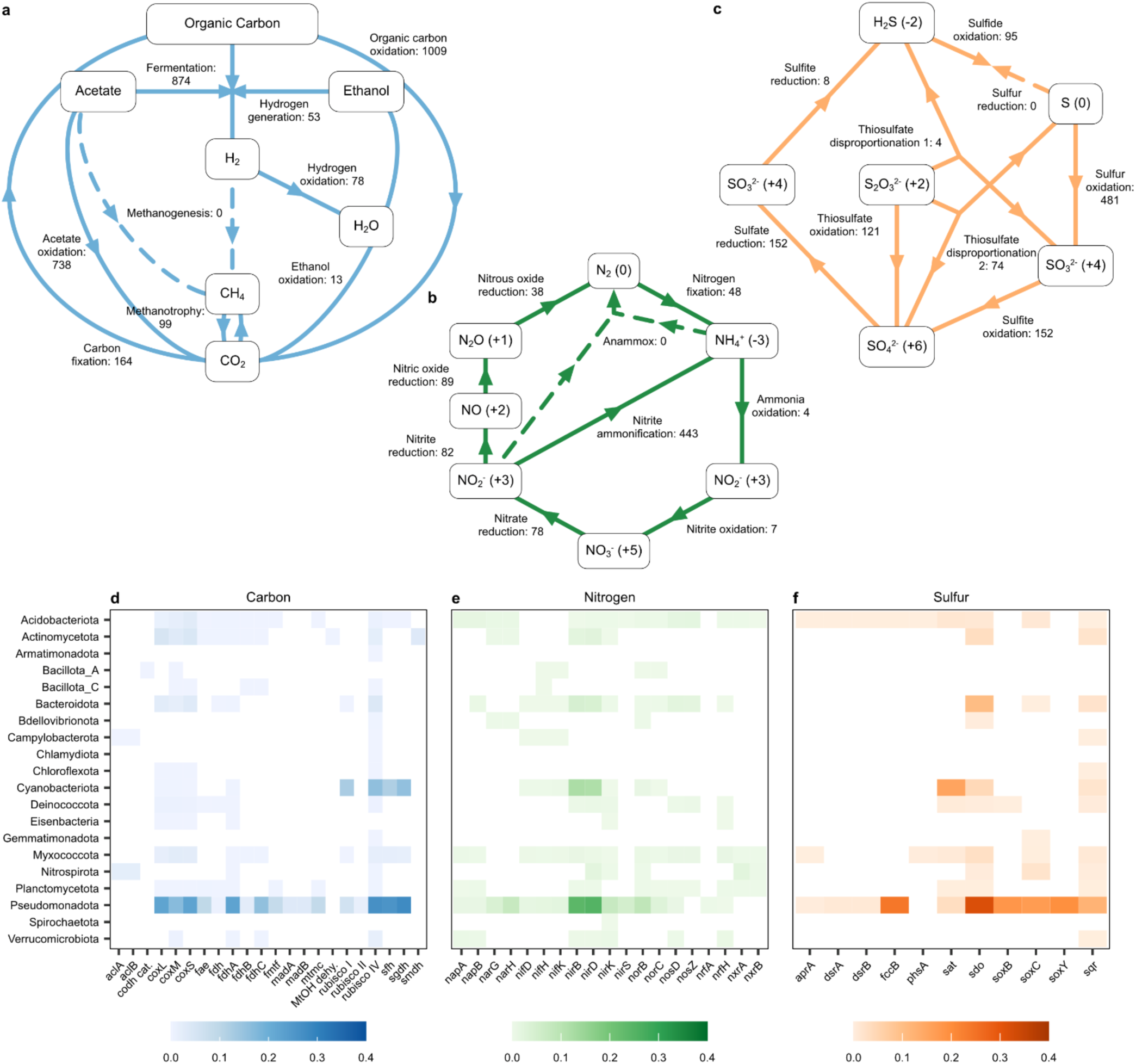
Nutrient cycling and metabolic potential of bacterial MAGs. **(a)** Carbon, **(b)** nitrogen, and **(c)** sulfur cycling pathways, where numbers denote the number of MAGs positive for genes involved in each step and dashed lines show steps with no positive MAGs. Heatmaps of **(d)** carbon, **(e)** nitrogen, and **(f)** sulfur cycling genes where colour scales represent gene counts weighted by mean relative abundance and summed by bacterial phylum.

Biofilm bacteria exhibited an extensive range of functional capabilities, including resource acquisition, resource use, and stress tolerance (Fig. 5). Genes for the breakdown of compounds such as simple and complex carbohydrates, cellulose, and proteins were widely distributed across the community (present in between 75.83 and 100% of MAGs) (Fig. 5A). This indicates that the biofilm community can utilise a broad range of organic compounds, highlighting their metabolic flexibility and resilience to fluctuating nutrient availability.

**Fig. 5.**
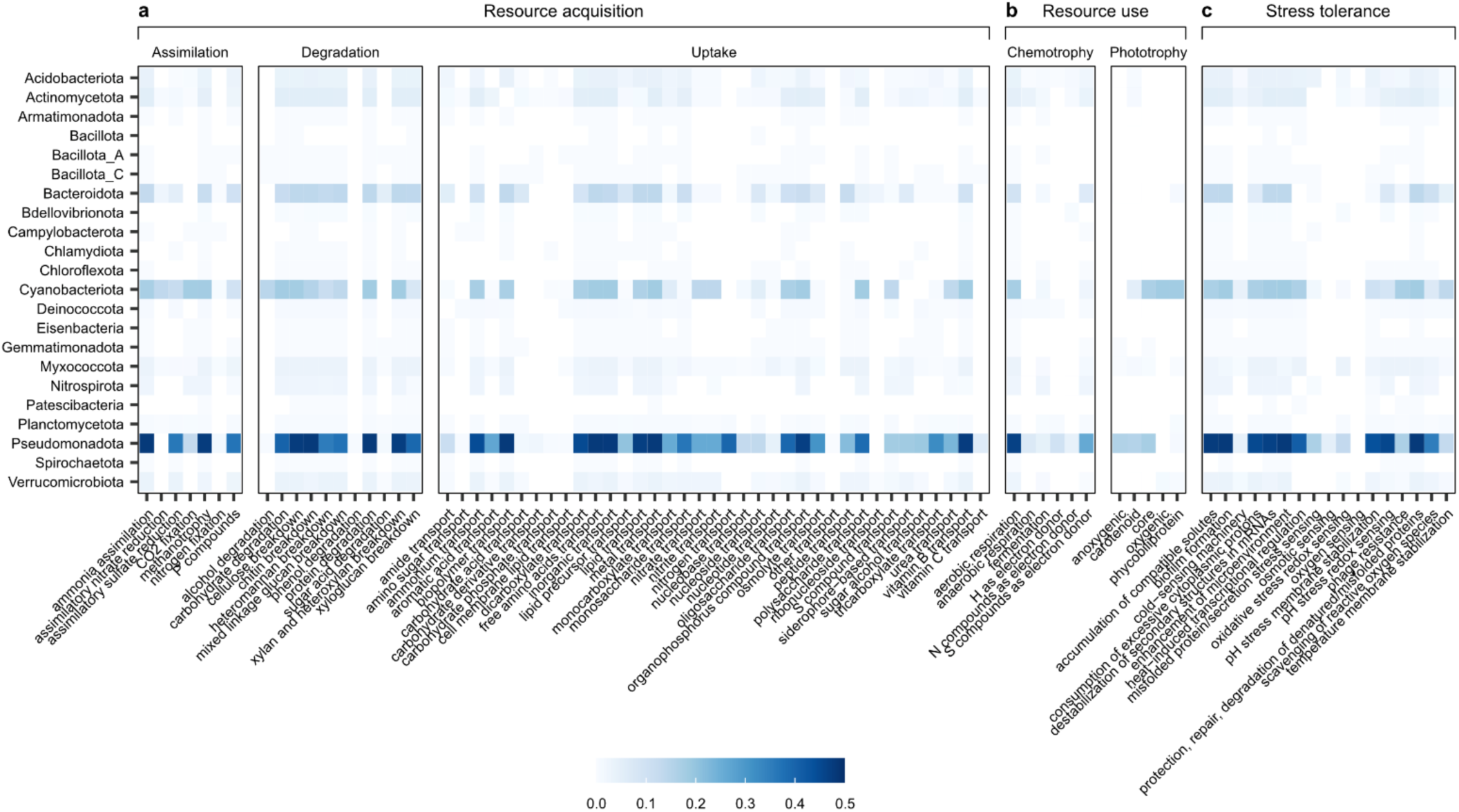
Functional genomic traits of bacterial MAGs. Traits associated with **(a)** resource acquisition (substrate assimilation, degradation, and uptake), **(b)** resource use (chemotrophy and phototrophy), and **(c)** stress tolerance. The heatmap colour scale represents gene counts per category weighted by mean relative abundance and summed by bacterial phylum.

Up to 90.63% of biofilm MAGs contained genes for aerobic respiration, and 21.89% of MAGs, which included members of Pseudomonadota, Actinomycetota, Acidobacteriota, Deinococcota, and Planctomycetota, also possessed genes associated with anaerobic respiration. Additionally, some phyla, notably Pseudomonadota and Cyanobacteriota, had genes for both core and accessory photosynthetic pigments (Fig. 5B). The diverse trophic capabilities identified among biofilm bacteria likely reflect niche adaptation to the stratified microenvironment within the biofilm. In this structure, aerobic and photosynthetic bacteria dominate the outer layer of the biofilm that is exposed to light and oxygen, whereas anaerobes occupy deeper oxygen-depleted layers. This stratification facilitates resource flow within the biofilm, where byproducts from one group can serve as substrates for another, creating a highly interconnected community^9^.

Biofilm formation is a key survival strategy for many freshwater microbes, allowing them to accumulate and readily access nutrients, gain protection from the extracellular polymeric matrix, and interact with other taxa^9^. Genes associated with biofilm formation were identified in 97.04% of MAGs encompassing nearly all bacterial phyla detected (Fig. 5C). Exceptions to this were Patescibacteria, which are known for their reduced genomes and symbiotic lifestyle^41^ and likely depend on interactions with biofilm-forming bacteria rather than contributing to biofilm formation directly, and Bacillota, which may play a less prominent role in biofilm formation, possibly by relying on an alternative survival strategy such as sporulation^42^. These findings highlight the widespread importance of biofilm formation as a strategy for microbial survival in dynamic river ecosystems, promoting community stability, and facilitating interactions among taxa. Additionally, a variety of other stress tolerance strategies were identified in the biofilm community. These include the ability to accumulate compatible solutes; stabilise membranes; scavenge reactive oxygen species; and protect, repair, and degrade damaged proteins (Fig. 5C).

Biofilm communities play an essential role in the uptake, transformation, and degradation of pollutants that are harmful to river ecosystems^39^. Most bacterial phyla (and 92.11% MAGs) contained genes involved in organophosphorus transport (Fig. 5A), which is a common pollutant in rivers derived from agricultural pesticides^43^. Members of Pseudomonadota, Actinomycetota, Deinococcota, Myxococcota, Cyanobacteriota, and Verrucomicrobia (and 21.70% of MAGs) had genes for the transport of aromatic acids such as polycyclic aromatic hydrocarbons (PAHs), which are harmful compounds that are introduced into rivers from industrial activities^44^. Furthermore, all bacterial phyla (and 99.51% of MAGs) had genes involved in the transport of metals, which are released into rivers from industrial discharge, mining, and other anthropogenic activities^45^. The widespread presence of these genes indicate that biofilm bacteria are capable of transporting and therefore potentially transforming harmful organic pollutants that pose a significant risk to aquatic life in rivers. Biofilm bacteria may contribute to metal detoxification by immobilising metals and reducing their bioavailability in the water. This demonstrates the importance of biofilm bacteria in maintaining water quality^10^ and their valuable application as bioindicators for monitoring ecosystem health^46^, particularly as rivers are under increasing threat from pollution and contaminants worldwide^3^.

### Environmental drivers of river biofilm communities

Our findings demonstrate the taxonomic diversity of biofilm bacteria and their extensive metabolic and functional roles which are essential for riverine ecosystem functioning. However, while previous studies have explored the environmental drivers of river biofilm communities at the individual catchment scale^13,16,47^, large-scale patterns and drivers remain poorly understood^25^, particularly in comparison to surface water communities^4^. This knowledge gap limits our ability to understand and predict how biofilms and wider river ecosystems respond to environmental change across diverse landscapes. By incorporating detailed spatial information of the upstream catchment alongside high-resolution monitoring of water chemistry, this study represents a novel and comprehensive effort to assess the environmental drivers of river biofilm bacteria across large spatial scales.

Variance partitioning analysis was used to identify the environmental factors shaping river biofilm community composition across England. Environmental variables were categorised as catchment land cover, catchment geology, water chemistry and watershed characteristics. Collectively, these environmental factors explained between 52.33 (Bdellovibrionota) and 91.88% (Deinococcota) of variation in biofilm community composition at the phylum level (Fig. 6A). Catchment land cover has previously been shown to significantly influence bacterial community composition in New Zealand stream biofilms^19,25^, and in North American river surface water^4^. Consistent with these findings, upstream catchment land cover was also identified as an important driver of river biofilm communities in England, accounting for between 2.24 and 29.27% of observed variance, with the dominant land cover types varying among the bacterial phyla (Fig. 6C).

**Fig. 6.**
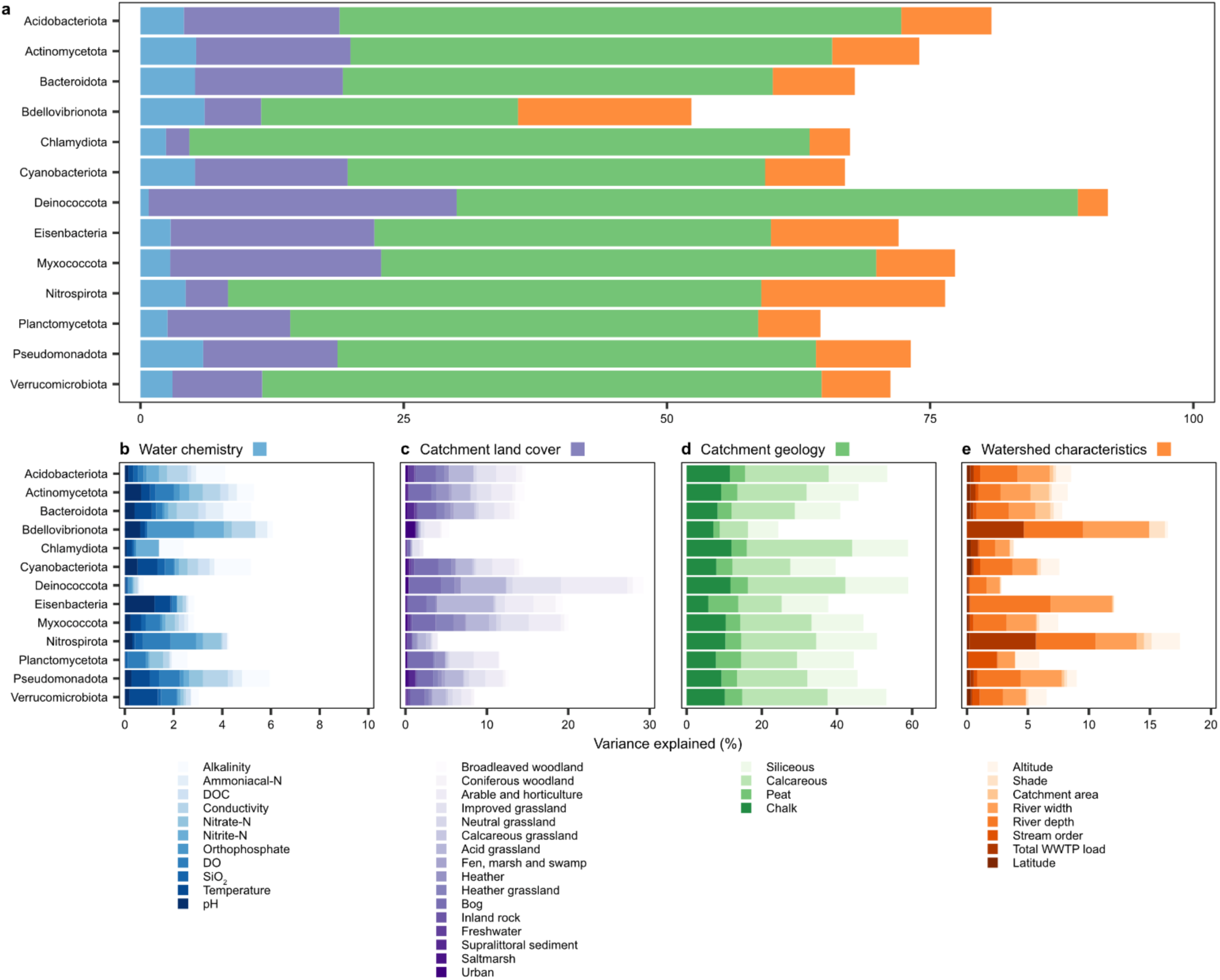
Environmental drivers of river biofilm bacterial communities. **(a)** Total variance explained by water chemistry, upstream catchment land cover, upstream catchment geology, and watershed characteristics, and variance explained by **(b)** water chemistry variables, **(c)** upstream catchment land cover types, **(d)** upstream catchment geology types, and **(e)** watershed characteristics. Variance explained for each bacterial phylum is calculated as a mean of variance explained at the individual MAG level.

Upstream catchment geology was found to account for the largest proportion of variance in biofilm bacterial community composition, explaining between 24.39 to 58.99% across bacterial phyla. The proportion of the upstream catchment represented by calcareous and siliceous geologies were identified as the dominant geological drivers, explaining up to 28.05 and 16.73% of community variation, respectively (Fig. 6D). Calcareous geology correlated positively with surface water pH (r = 0.24, p <0.001), alkalinity (r = 0.33, p <0.001), and conductivity (r = 0.24, p <0.001), while siliceous geology correlated negatively with these variables (r = −0.17, r = −0.53, and r = −0.26, respectively, p <0.001) (Extended Data Fig. 5, Supplementary Data 5). The influence of geology on biofilm communities therefore likely operates through its effects on pH, alkalinity, and conductivity^38,48^.

Water chemistry explained a relatively small proportion of variance (0.77 to 6.08%) (Fig. 6B). The physiochemical conditions of the surface water may be less ecologically relevant to the biofilm community compared to the geological properties and mineral composition of the stones on which biofilms develop and the microenvironment within the biofilm itself, which may be buffered against physicochemical fluctuations in the water column^9^. However, factors such as alkalinity, conductivity, pH, dissolved oxygen, and surface water temperature were still influential, each explaining up to 1.5% of the observed variation in any bacterial phylum. Furthermore, orthophosphate, nitrate-N, and nitrite-N emerged as key nutrients shaping biofilm community composition, most notably influencing the relative abundance of Nitrospirota (1.05, 0.76, and 0.27%, respectively) and Bdellovibrionota (1.92, 0.32, and 1.21%, respectively). Wastewater treatment plant (WWTP) load which correlated positively with orthophosphate (r = 0.65, p <0.001), nitrate-N (r = 0.48, p <0.001) and nitrite-N (r = 0.56, p <0.001) (Extended Data Fig. 5), was also identified as a key driver of Nitrospirota and Bdellovibrionota relative abundance, accounting for 5.46 and 4.66% of variance, respectively (Fig. 6E). These results suggest that WWTP effluent which introduces organic nutrients into the watercourse can impact the dynamics of biofilm bacterial communities with implications for water quality and microbial nutrient cycling^17^.

The River Continuum Concept (RCC)^49^ describes how continuous physical gradients along a river’s course, from headwaters to mouth, shape its biological and chemical properties. Low-order streams are shallow, narrow and strongly influenced by shading and allochthonous input from vegetation. As stream order, and therefore river width and depth, increase towards the mouth, autochthonous inputs play a progressively larger role^49^. Successional shifts in surface water bacterial communities along the river continuum have been well-documented within individual catchments^50,51,52^. This trend extends to the national-scale, where stream order was identified as the most significant environmental driver shaping bacterial communities in the surface water of rivers across North America^4^, ranging from stream order 1 to 12. However, stream order within the RSN, which ranged from 1 to 8, was not a dominant driver of bacterial community composition in river biofilms across England, explaining less than 0.63% of variance for each bacterial phylum with the exception of Planctomycetota (2.38%) (Fig. 6E). Associated physical factors such as river depth and width explained up to 6.57 and 5.42%, respectively, but shading accounted for less than 1.28% of bacterial community composition. Bacterial communities within stream biofilms from a pre-alpine catchment were also found to show patterns contradictory to the RCC^53^. These contrasting findings between river surface water and biofilm bacterial communities provide evidence that successional changes along the river continuum affect these distinct microbial communities differently. While variation in surface water bacterial communities can be primarily explained by the RCC, local factors such as geology, land cover, nutrients and WWTP load are more influential in shaping river biofilm communities. Furthermore, cross-domain biotic interactions may also be an important driver of biofilm bacterial community dynamics and require further study^53^.

### Conclusion

Rivers are valuable, biodiverse habitats and their microbial communities, particularly those within biofilms, are critical to ecosystem functioning, biogeochemical cycling, pollutant degradation, and maintaining water quality^1,2^. By leveraging comprehensive national-scale sampling coupled with high-resolution environmental data, this study provides novel insights into the biogeography, taxonomic diversity, metabolic potential, and functional roles of river biofilm microbial assemblages.

River biofilms are revealed to be dominated by high-occupancy generalist bacteria, most notably members of Pseudomonadota, Cyanobacteriota and Bacteroidota, reflecting the interconnectedness of river ecosystems that facilitate microbial dispersal across the river network and colonisation by generalist taxa. These results further highlight the significant metabolic and functional versatility of river biofilms, which harbour a diverse array of genes supporting carbon, nitrogen and sulfur cycling, the utilisation of a broad range of organic compounds, the transportation of organic pollutants, and various trophic strategies. Their capacity to perform diverse biochemical roles reflects niche adaptation to the biofilm microenvironment and provides resilience in dynamic river ecosystems, allowing them to sustain key biogeochemical processes under fluctuating or unfavourable conditions.

The RCC is a strong driver of surface water bacterial communities^4,50,51,52^. However, this framework is less applicable to river biofilm communities. Instead, upstream catchment geology and land cover emerge as the primary determinants of biofilm communities, which likely shape community composition through their effects on the physicochemical conditions of the water column and importantly the biofilm microenvironment. Furthermore, anthropogenic activities, including changes in land use and the release of WWTP effluents and pollutants into rivers, may significantly impact biofilm community dynamics. River biofilms are valuable bioindicators of ecosystem health^46^, and our findings have the potential to inform biodiversity conservation practices, water quality monitoring and management, and public health measures, and, importantly, explain resilience to environmental change observed in river biofilm bacterial communities.

## Methods

### Sample collection

A total of 450 river biofilm samples were collected over a three-year period (2021-2023) from 146 sites within the EA’s RSN programme. Biofilms were collected from the river benthos by scraping biofilm-covered stones from the riverbed at each site with a sterile toothbrush and deionised water^54^. At sites where stones were not accessible, biofilms were collected by macrophyte scrape. The suspended biofilm was transferred to a 15 ml tube and preserved in the field in 5 ml DNA preservation buffer (3.5 M ammonium sulphate, 17 mM sodium citrate and 13 mM EDTA). Samples were immediately transported to the EA’s National Laboratory at Starcross, Exeter where they were concentrated by centrifugation and frozen. Samples were then transported to the UK Centre for Ecology & Hydrology (UKCEH), Wallingford on dry ice and stored at −20 °C.

### Water chemistry

Water temperature (°C), pH, alkalinity to pH 4.5 as CaCO_3_ (mg L^-1^), conductivity, and the concentration of dissolved oxygen (DO, mg L^-1^), dissolved organic carbon (DOC, mg L^-1^), orthophosphate (mg L^-1^), nitrate-nitrogen (nitrate-N, mg L^-1^), nitrite-nitrogen (nitrite-N, mg L^-1^), ammoniacal nitrogen (ammonia-N, mg L^-1^), and reactive SiO_2_ (mg L^-1^) were measured using surface water samples collected from each sampling site and this data is available at: https://environment.data.gov.uk/water-quality/view/landing. For each variable, a mean was calculated from up to 5 independent measurements taken during a 3-month period prior to biofilm sampling.

### Upstream catchments

The upstream catchment of each RSN site was determined using 10 m flow direction and accumulation grids and the sampling points which were snapped to the maximum flow accumulation cell within 50 m of its location. The digital elevation model has a 10 m resolution and was based on NEXTMap which was derived from airborne Interferometric Synthetic Aperture Radar (IFSAR) and is available on the UKCEH Catchment Management Modelling Platform (https://catalogue.ceh.ac.uk/cmp/documents). The area of the derived shapefile was then used as the area of the upstream catchment (km^2^). The upstream catchment shapefiles were intersected with the 10 m scale 2020 and 2021 UKCEH land cover maps available at: https://www.ceh.ac.uk/data/ukceh-land-cover-maps, and the British Geological Survey (BGS) geology map available at: https://www.bgs.ac.uk/datasets/bgs-geology-250k. The percentage of the upstream catchment area covered by each land cover type or geology was calculated for each sampling site.

### Watershed characteristics

Strahler numbers for the branches of the rivers where sampling sites are located were determined by spatially joining the GRTS points with the 2021 OS Open Rivers Network layer (https://www.ordnancesurvey.co.uk/products/os-open-rivers). To provide an estimate of river depth and width at sampling sites, sampling points were intersected with the gridded 1 km^2^ physical river characteristics for the UK v2 dataset (https://doi.org/10.5285/8df65124-68e9-4c68-8659-1c6b82c735e9).

To determine the number of wastewater treatment plants (WWTPs) in the upstream catchments of each site and the total population equivalent loads, a spatial intersect was performed between the upstream catchments and a dataset of WWTPs in England and Wales with their population equivalent loads mapped to discharge location (https://www.data.gov.uk/dataset/0f76a1c3-1368-476b-a4df-7ef32bfd9a8b/urban-waste-water-treatment-directive-treatment-plants).

Count and sum functions were used to provide the number of WWTPs and their total population equivalent loads respectively.

Average shading from landforms and surface model objects was determined for each sampling by spatially joining the sampling points with a 25 m buffer zone to the Environment Agency Keeping Rivers Cool relative shading map available at: https://data.catchmentbasedapproach.org/maps/theriverstrust::riparian-shade-england.

### DNA extraction

Each biofilm sample was defrosted and briefly vortexed to homogenise the sample. The Quick-DNA fecal/soil microbe kit (Zymo Research, California, U.S.) was used to extract DNA from 100 µL of sample with the following amendments to the manufacturer’s protocol to maximise DNA yield: Zymo DNA/RNA shield was used as the lysis buffer, samples were lysed at 20 Hz for 20 min using the TissueLyser II (Qiagen, Germany), and 20 µL of proteinase K was added to the lysate prior to incubation at 65 °C for 20 min. Purified DNA was eluted in 100 µL of elution buffer. The purity of extracted DNA was checked using the NanoDrop 8000 spectrophotometer (Thermo Fisher Scientific, MA, U.S.). The concentration of DNA was measured using the QuantiFluor ONE dsDNA kit (Promega, Madison, WI, U.S.) with a BioTek Cytation 5 imaging reader (Agilent Technologies, California, U.S.) according to the manufacturer’s protocol. The DNA was stored frozen at −70 °C for long-term archiving at UKCEH, Wallingford.

### Metagenomic sequencing

Extracted DNA was sent to Novogene UK for library preparation and 2x 150 bp shotgun metagenomic sequencing on an Illumina NovaSeq 6000 with an S4 flow cell to achieve a sequencing depth of at least 10 Gb raw data per sample. Between 2.85 million and 188.26 million raw reads were generated per sample, with an average of 66.80 million raw reads per sample.

### Metagenomic data processing

Illumina adaptor sequences were trimmed, and metagenomic reads were filtered to a minimum quality score of 25 using Trim Galore v0.6.5 (https://github.com/FelixKrueger/TrimGalore).

Reads mapping to the human reference genome (GRCh38; downloaded on 2023-11-16 from https://ftp.ncbi.nlm.nih.gov/refseq/H_sapiens/annotation/GRCh38_latest/refseq_identifiers/GRCh38_latest_genomic.fna.gz) were removed. After quality filtering and trimming, between 4.28 million and 90.17 million reads were obtained per sample, with an average of 37.41 million reads per sample. To determine the overall profile and percentage of archaea, bacteria, and eukaryotes in the dataset, singleM v0.16.0^55^ was run on the preprocessed reads. MultiQC v1.17^56^ was used to process all filtered reads to obtain basic statistics per sample. Megahit v1.2.9^57^ was used to assemble the reads into contigs, which were subsequently used for binning, functional annotation, and downstream analyses. For functional analysis, open-reading frames (ORFs) were predicted using Prodigal v2.6.3^58^, which were fed into EggNOG-mapper v2.1.9^59^ for functional annotations via the built-in v6.0 database. The ORFs were used in featureCounts^60^ to generate per-gene coverage, while Kraken2 v2.1.2^61^ and Bracken v2.6.0^62^ were used to taxonomically annotate the contigs, using the PlusPFP (https://benlangmead.github.io/aws-indexes/k2) database, which includes RefSeq protozoa, fungi, plant, archaea, bacteria, virus, plasmid, and human sequences. For stringency, a 0.7 confidence threshold was used in Kraken2. The contigs were also used to obtain metagenome-assembled genomes (MAGs), by dereplicating bins obtained via MetaBAT v2.15^63^, MetaBinner v1.4.3^64^ and CONCOCT v1.1.0^65^. The completion and contamination of the dereplicated bins were estimated with CheckM2 v1.2.2^66^, alongside taxonomic annotation using the GTDBtk v2.3.2^67^. Simultaneously, microTrait v1.0.0^68^ was used to extract fitness traits from the MAGs, based on protein family sequence similarities and Kyoto Encyclopedia of Genes and Genomes (KEGG) orthologs. The metabolic and biogeochemical traits of the MAGs were also analysed using METABOLIC v4.0^69^ and the metabolic pathways were further assessed via metabolisHMM v2.22^70^. The workflow was implemented using the Snakemake workflow management system v7.8.2^71^, and is available at: https://github.com/amycthorpe/metag_analysis_EA for preprocessing the metagenomic reads to MAG assembly. The analysis workflow, including dereplication, taxonomy assignment and the implementation of microTrait, METABOLIC and metabolisHMM, is available at: https://github.com/amycthorpe/EA_metag_post_analysis.

### Data analysis

Using contamination and completeness statistics estimated with CheckM2, the MAGs were categorised as near-complete (>90% complete and <5% contamination) or medium-quality (>70% complete and <10% contamination), with remaining MAGs that did not meet the thresholds categorised as low-quality. A phylogenetic tree was constructed using the maximum likelihood method with the ggtree R package v3.12.0^72^ using the multiple sequence alignment from the GTDB toolkit analysis. For biogeographic mapping, relative abundance was determined for each MAG, summed at the phylum level and plotted according to site latitude and longitude. For occupancy and relative abundance analyses, mean relative abundance was calculated for each MAG across all samples, and the number of samples each MAG was present in was used as a measure of occupancy. Levins’ niche breadth (Bn) was calculated for each MAG using the MicroNiche R package v1.0.0^73^. All MAGs were above the recommended limit of quantification threshold of 1.65. MAGs with a Bn greater than the median Bn (0.03) were categorised as generalists, and those with a Bn less than the median were categorised as specialists. The percentage of generalists and specialist MAGs was calculated for each bacterial phylum. The presence of metabolic and functional traits identified for each MAG using METABOLIC, metabolisHMM, and microTrait were weighted by mean relative abundance and summed at the phylum level.

Variance partitioning was used to identify the environmental drivers of bacterial community composition at the MAG level. A linear mixed model with adjustment for multiple comparisons was fitted to the matrix of MAG relative abundance to estimate the contribution of each environmental variable to community variation using the variancePartition R package v1.34.0^74^. Samples (n=450) were filtered to complete observations according to environmental data availability (n=401). Mean variance explained was then calculated for each bacterial phylum. Due to low variance for some MAGs, variance partioning was based on a subset of 335 MAGs. Pearson correlations (two-sided) were computed using the cor() function in R to investigate correlation between the environmental variables and bacterial phyla on the subset of 401 samples with complete observations. Data analysis was performed in R v4.4.0^75^, and R scripts for data analysis and visualisation are available at: https://github.com/amycthorpe/biofilm_MAG_analysis.

## Data availability

Metagenomic reads have been deposited in the European Nucleotide Archive (ENA) at EMBL-EBI under accession number PRJEB85861. Sample accession codes, the assembled MAGs, and all the data presented, including the environmental metadata associated with each sample, MAG coverage, taxonomy, and CheckM2 statistics, outputs from METABOLIC, metabolisHMM, and microTrait, and the niche breadth index, variance partitioning, and correlation analysis results are available on Zenodo at: https://doi.org/10.5281/zenodo.14864644.

## Code availability

Snakemake workflows used to process and analyse the metagenomic data are available at: https://github.com/amycthorpe/metag_analysis_EA and https://github.com/amycthorpe/EA_metag_post_analysis. R scripts for data analysis and visualisation are available at: https://github.com/amycthorpe/biofilm_MAG_analysis.

## Acknowledgements

Thank you to Dr Jonathan Porter, Sean Butler, and Alan Wan from the Environment Agencies National Monitoring Laboratories for their support collating and shipping the biofilm samples for further processing. This work was funded by the Environment Agency under research project SC220034. ACT, SBB and DSR were supported by Natural Environment Research Council (NERC) grant NE/X015947/1. KW and JW were supported by NERC grant NE/X015777/1. The authors acknowledge the support of the Biotechnology and Biological Sciences Research Council (BBSRC), part of UK Research and Innovation; Earlham Institute Strategic Programme Grant Decoding Biodiversity BB/X011089/1 and its constituent work package - BBS/E/ER/230002C (Decode WP3 Linking Fine-Scale Microbial Diversity to Ecosystem Functions).

## Author contributions

**ACT**: conceptualisation, data curation, investigation, formal analysis, visualisation, writing – original draft. **SBB**: conceptualisation, data curation, formal analysis, writing – review and editing. **JW**: conceptualisation, data curation, writing – review and editing. **LHH**: data curation, writing – review and editing. **KW**: conceptualisation, supervision, writing – review and editing. **DSR**: conceptualisation, supervision, writing – review and editing.

## Corresponding authors

Correspondence should be addressed to Amy C. Thorpe, Susheel Bhanu Busi or Daniel S. Read.

## Competing interest declaration

The authors declare no competing interests.

## Supplementary Information

**Supplementary Data 1**

environmental_metadata.xlsx: environmental metadata associated with each sample, sheet 1: water chemistry, sheet 2: catchment land cover, sheet 3: catchment geology, sheet 4: watershed characteristics, sheet 5: data sources.

**Supplementary Data 2**

singlem_results.csv: proportion of archaea, bacteria, and eukaryotes in the preprocessed metagenomic reads.

**Supplementary Data 3**

MAG_metadata.xlsx: metadata associated with bacterial MAGs, sheet 1: coverage, sheet 2: CheckM2 statistics, sheet 3: GTDB taxonomy, sheet 4: Levins’ index.

**Supplementary Data 4**

metabolism_and_functions.xlsx: presence of metabolic and functional traits identified in the bacterial MAGs, sheet 1: METABOLIC results, sheet 2: metabolisHMM results, sheet 3: microTrait results.

**Supplementary Data 5**

varPart_and_correlations.xlsx: variance partitioning and correlation analysis, sheet 1: variance partitioning, sheet 2: correlations.

## Extended Data

**Extended Data Table 1.** Number of sampling sites and the dominant land cover type and geology in the upstream catchment.

| Dataset | Type | Number of sites |
| --- | --- | --- |
| Land use | Acid grassland | 24 |
|  | Arable and horticulture | 47 |
|  | Bog | 5 |
|  | Broadleaved woodland | 2 |
|  | Coniferous woodland | 2 |
|  | Heather | 5 |
|  | Heather grassland | 4 |
|  | Improved grassland | 54 |
|  | Neutral grassland | 1 |
|  | Suburban | 2 |
| Geology | Calcareous | 77 |
|  | Chalk | 19 |
|  | Peat | 7 |
|  | Siliceous | 43 |

**Extended Data Table 2.** Summary of water chemistry data.

|  | Mean | SD | SE | Min | Max | Median | Count |
| --- | --- | --- | --- | --- | --- | --- | --- |
| Alkalinity | 152.73 | 86.74 | 4.31 | 5.00 | 446.67 | 163.33 | 405 |
| Ammoniacal-N | 0.08 | 0.25 | 0.01 | 0.00 | 4.57 | 0.04 | 443 |
| Dissolved organic C | 4.93 | 4.13 | 0.20 | 0.33 | 41.50 | 3.75 | 443 |
| Conductivity | 521.13 | 553.03 | 26.25 | 24.00 | 7748.33 | 458.33 | 444 |
| Nitrate-N | 4.06 | 4.02 | 0.19 | 0.00 | 20.57 | 2.79 | 443 |
| Nitrite-N | 0.02 | 0.03 | 0.00 | 0.00 | 0.26 | 0.01 | 443 |
| Orthophosphate | 0.11 | 0.35 | 0.02 | 0.00 | 6.20 | 0.04 | 443 |
| Dissolved oxygen | 10.70 | 1.87 | 0.09 | 2.55 | 14.70 | 10.88 | 444 |
| SiO<sub>2</sub> | 7.07 | 3.44 | 0.16 | 1.02 | 24.00 | 6.80 | 441 |
| Water temperature | 11.02 | 4.13 | 0.20 | 2.30 | 19.97 | 11.50 | 444 |
| pH | 7.82 | 0.53 | 0.02 | 5.18 | 8.80 | 7.93 | 444 |

Mean, standard deviation (SD), standard error (SE), minimum, maximum, and median are calculated from the 3 months mean (mean of up to 5 measurements taken during a 3-month period prior to biofilm sampling) across up to 450 samples collected from 146 sites between 2021 and 2023. Count is the number of river biofilm samples with a paired water chemistry measurement.

**Extended Data Fig. 1.**
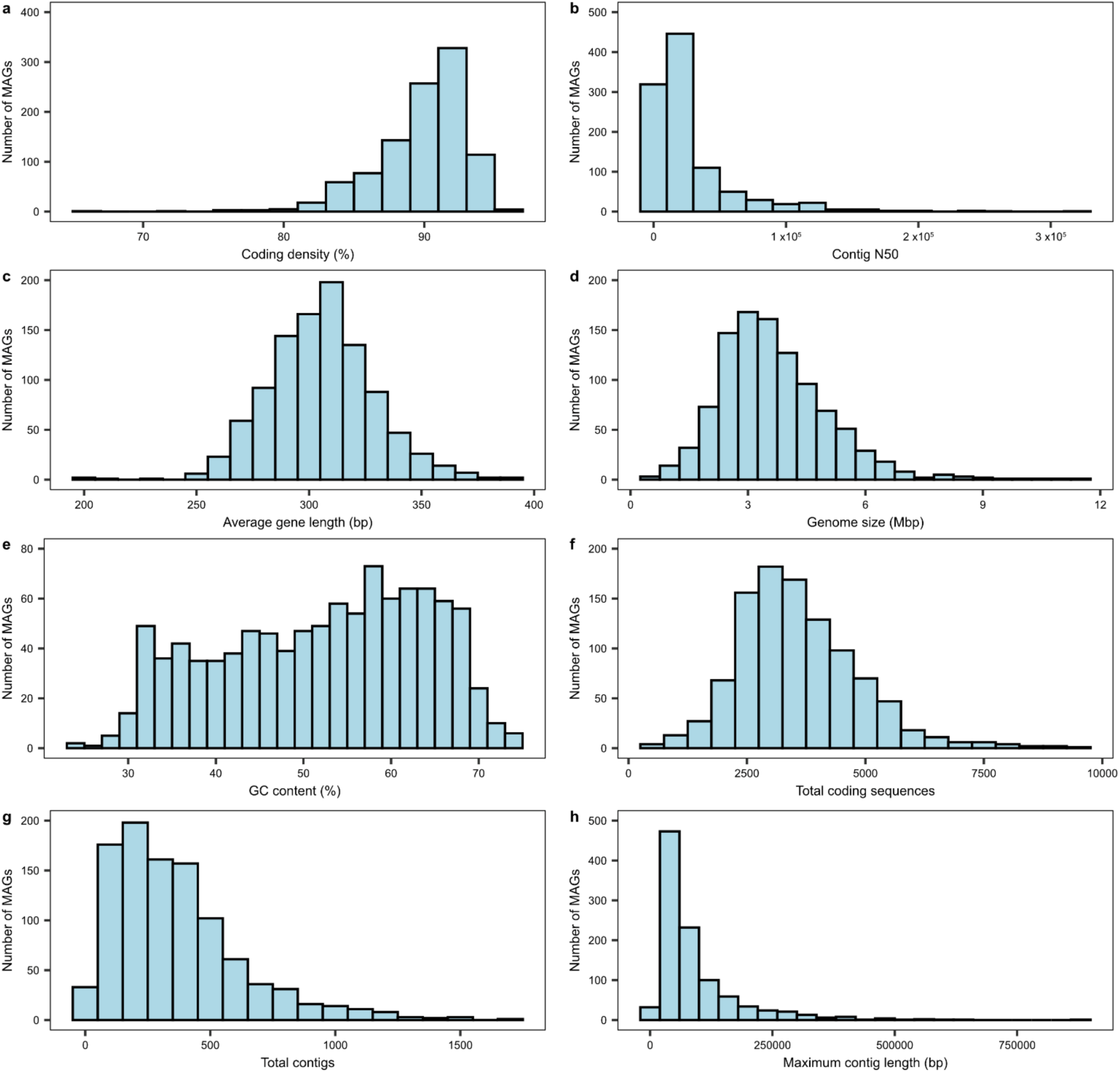
Distribution of MAG traits. **(a)** Coding density, **(b)** contig N50, **(c)** average gene length, **(d)** genome size, **(e)** GC content, **(f)** total coding sequences, **(g)** total contigs, and **(h)** maximum contig length.

**Extended Data Fig. 2.**
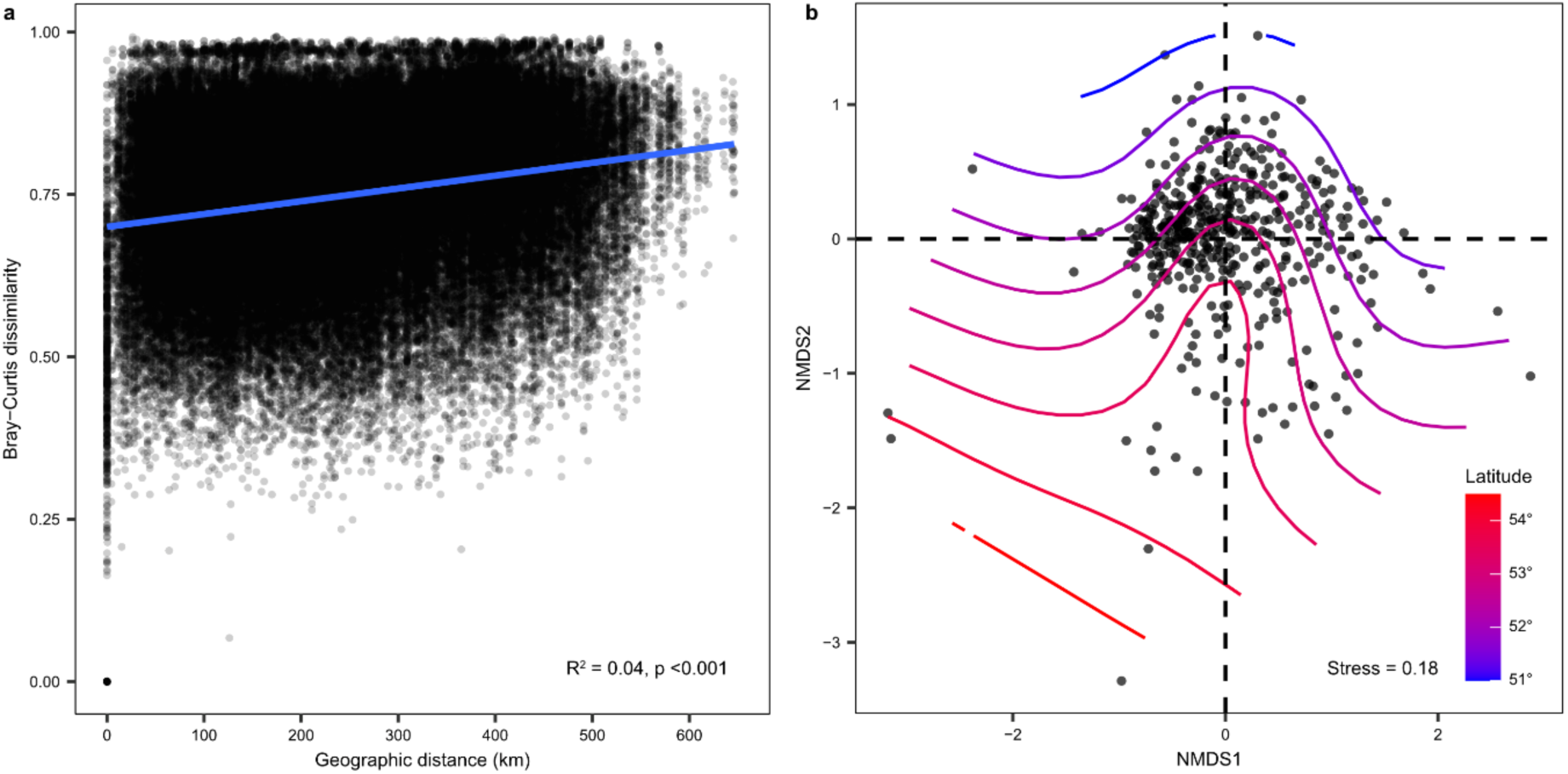
Distance-decay and beta-diversity of river biofilm bacterial communities. **(a)** Distance-decay relationship between Bray-Curtis community dissimilarity and geographic distance, R^2^ and p values of the linear regression are shown. **(b)** Non-metric multidimensional scaling (NMDS) of a Bray-Curtis dissimilarity matrix based on beta diversity, latitude is represented by contour lines where the red to blue gradient indicate north to south.

**Extended Data Fig. 3.**
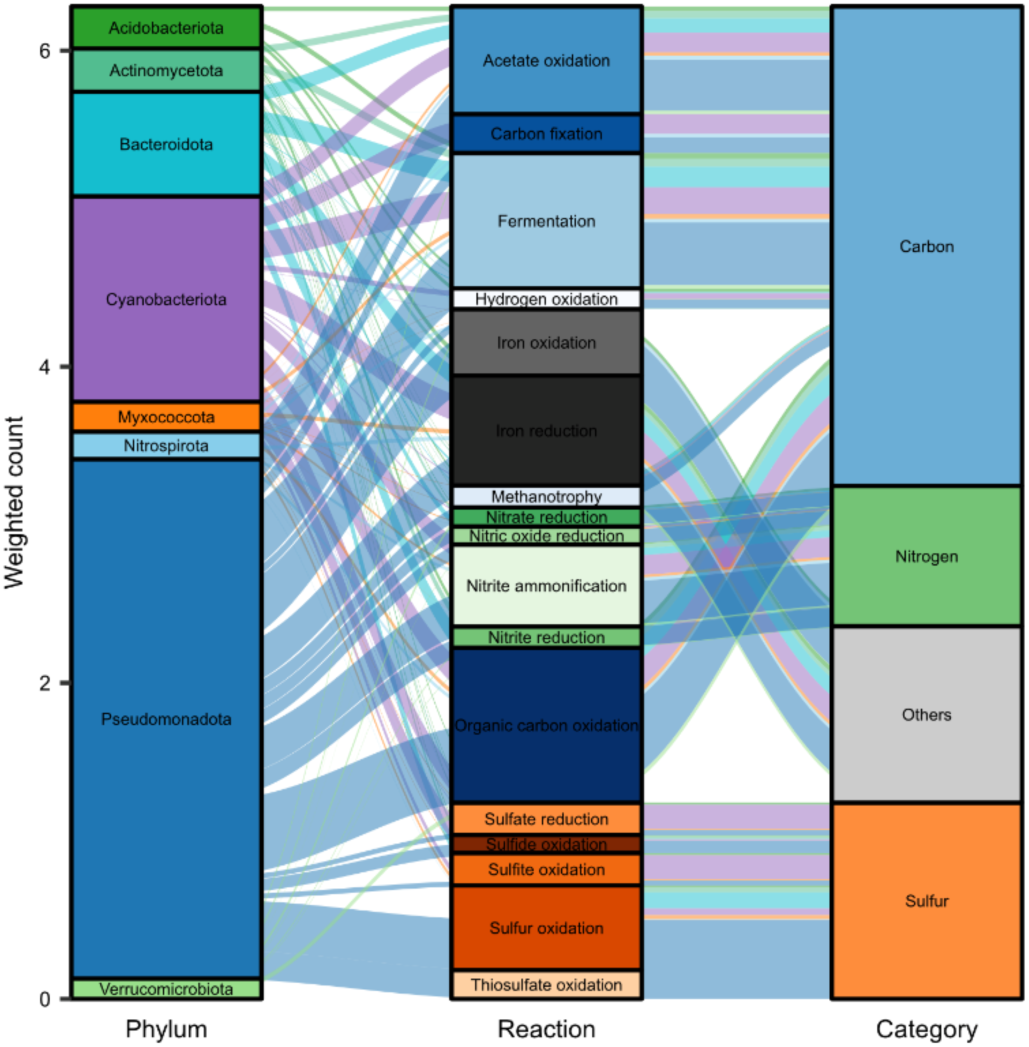
Alluvial plot of metabolic pathways identified in the MAGs. Weighted count is gene count weighted by mean relative abundance and summed by bacterial phylum. Only phyla and reactions with total weighted count >0.1 are shown for clarity.

**Extended Data Fig. 4.**
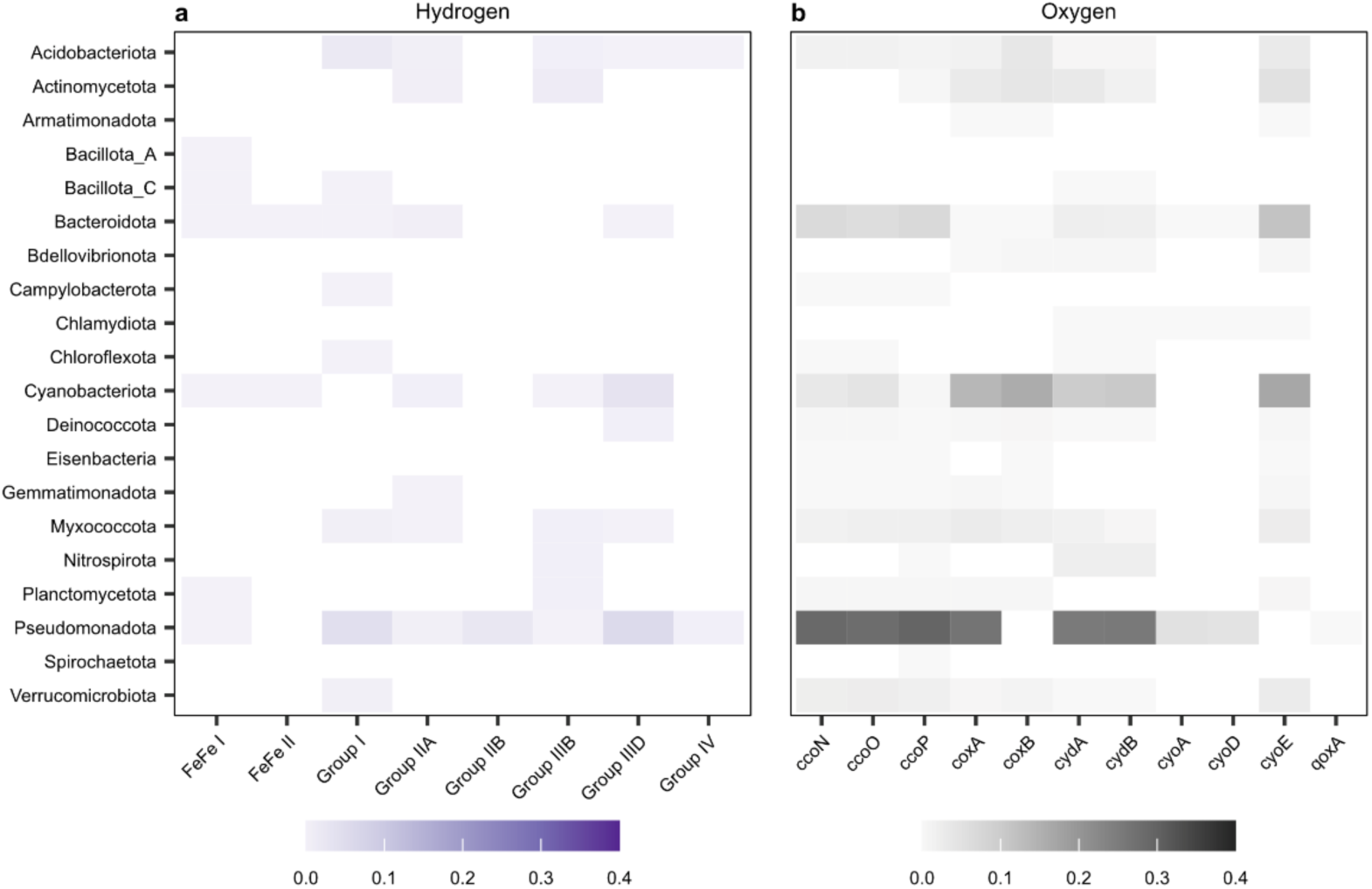
Hydrogen and oxygen cycling potential of bacterial MAGs. Heatmaps of **(a)** hydrogen, and **(b)** oxygen cycling genes where colour scales represent gene counts weighted by mean relative abundance and summed by bacterial phylum.

**Extended Data Fig. 5.**
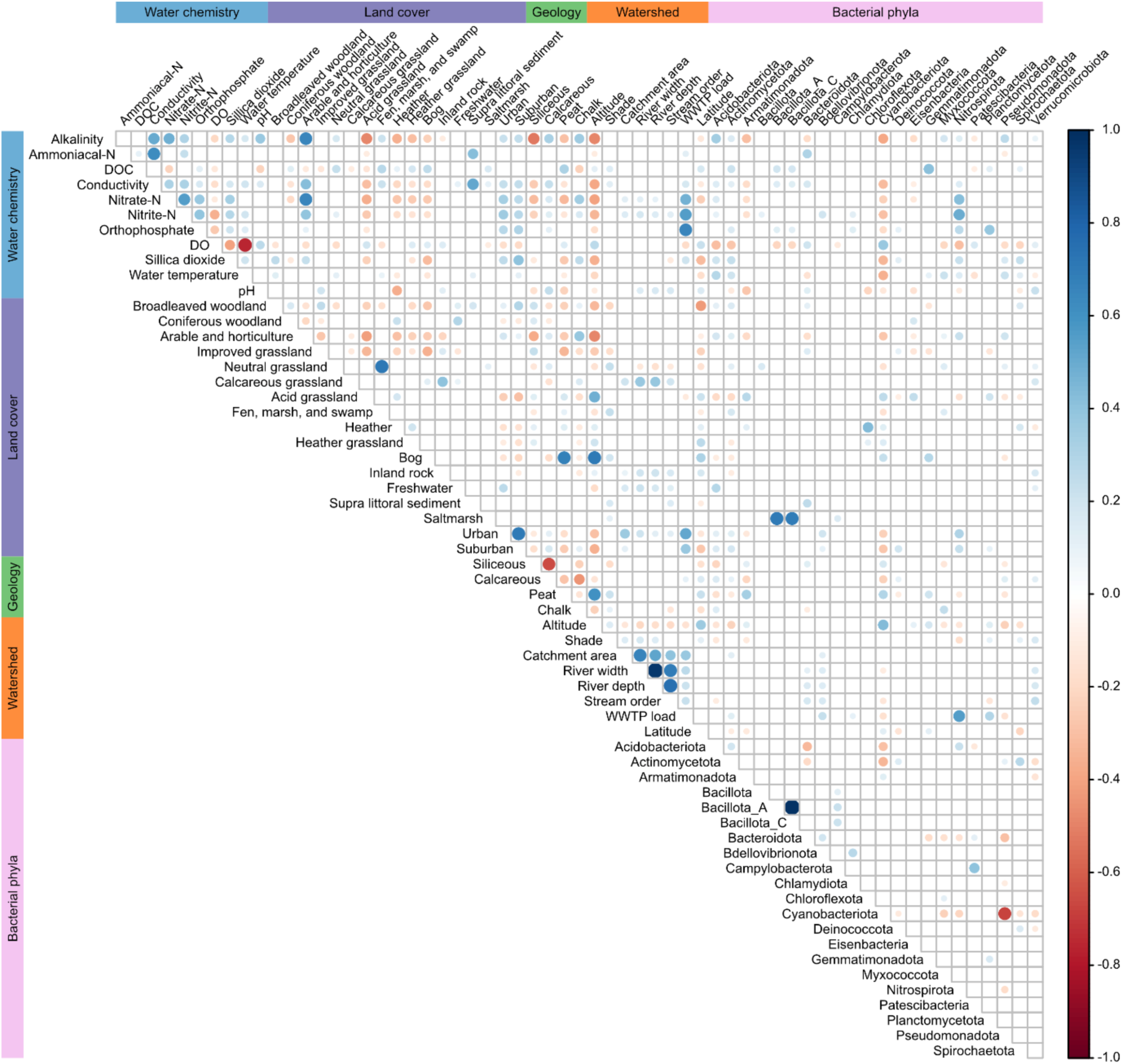
Pearson correlations between environmental variables and bacterial phyla. Heatmap scale indicates strength of correlation, with blue representing positive correlations and red representing negative correlations. Point size is proportional to the strength of correlation. Only significant (p <0.05) correlations are shown.

## Notes

https://zenodo.org/records/14864644

### Competing Interest Statement

The authors have declared no competing interest.

